# Concurrent maintenance of both veridical and transformed working memory representations within unique coding schemes

**DOI:** 10.1101/2022.10.28.514218

**Authors:** Güven Kandemir, Michael J. Wolff, Aytaç Karabay, Mark G. Stokes, Nikolai Axmacher, Elkan G. Akyürek

**Author notes:** shared first authorship (Güven Kandemir, Michael J. Wolff). Address correspondence to: Elkan Akyürek Department of Experimental Psychology, University of Groningen Grote Kruisstraat 2/1, 9712 TS Groningen, The Netherlands.

## Abstract

In the dynamic environment we live in, the already limited information that human working memory can maintain needs to be constantly updated to optimally guide behaviour. Indeed, previous studies showed that leading up to a response, representations maintained in working memory representations are transformed continuously. This goes hand-in-hand with the removal of task-irrelevant items. However, does such removal also include the representations of stimuli as they were originally, prior to transformation? Here we assessed the neural representation of task-relevant transformed representations, and the no-longer-relevant veridical representations they originated from. We applied multivariate pattern analysis to electroencephalographic data during maintenance of orientation gratings with and without mental rotation. During maintenance, we perturbed the representational network by means of a visual impulse stimulus, and were thus able to successfully decode veridical as well as imaginary, transformed orientation gratings from impulse-driven activity. The impulse response reflected only task-relevant (cued), but not task-irrelevant (uncued) items, suggesting that the latter were quickly discarded from working memory. By contrast, even though the original cued orientation gratings were also no longer task-relevant after mental rotation, these items continued to be represented next to the rotated ones, in different representational formats. This seemingly inefficient use of scarce working memory capacity was associated with reduced probe response times and may thus serve to increase precision and flexibility in guiding behaviour in dynamic environments.

## 1. Introduction

Working memory (WM) is more than a short-term storage for sensory input; it comprises the ability to actively manipulate and modify information as well (Baddeley, 2003). This is essential, since our environment is constantly in flux, and we need to organize perceptual information in that dynamic context. Indeed, one of the key features of WM is that it can adapt and alter its contents to anticipate change. For example, the egocentric location of an object will change constantly as we move around, yet we can easily point at its current spatial location even when it is temporarily out of sight (Rieser et al., 1986), showing that previous sensory input can be recalled and transformed to predict its changed state. Thus, the ability to not only maintain but also add information through imagery, and to update existing information in WM, is essential to guide both current and future behaviour (Rainer et al., 1999).

The brain seems to use the same neural substrate to maintain both current and previous sensory inputs, as well as mentally imagined items. Imagery and sensory-driven perception activate spatially overlapping regions in visual cortex, and share a common coding scheme (Stokes et al., 2009). Furthermore, items maintained in WM as well as items that are only imagined are both represented in early visual brain regions, with common activity patterns resembling those generated by direct visual input (Albers et al., 2013). These neural commonalities make a functional interpretation seem appealing (but see also Linke & Cusack, 2015; Iamshchinina et al., 2021), and several authors have highlighted the similarity of WM, imagery, and perception (Dijkstra et al., 2019; Pearson, 2019), all of which may use primarily sensory brain regions, such as visual cortex, as a “blackboard” (Roelfsema & De Lange, 2016).

Loaded with such wide-ranging demands, an important question is how the brain keeps its blackboard clean and fit for re-use. Not only that, but it also needs to be able to distinguish between current sensory input and maintained or imagined information. This is further compounded by the fact that it is not sufficient to retain only stimulus-related information, since WM ultimately serves to guide behaviour (Stokes et al, 2013; van Ede et al., 2019). In many cases, information that was once encoded to reflect certain properties of the environment may thus have to be updated or even superseded to relate to a possible course of action. On the one hand, it could be an economical strategy to free up WM capacity by removing or overwriting information that is no longer behaviourally relevant. On the other hand, preserving such information might carry the adaptive advantage that transformations could be re-applied on their originals, or even adjusted if the need arose. To arbitrate between these alternatives was the purpose of the present study.

There is at least some prior evidence from fMRI to suggest that once an item is transformed (e.g., mentally rotated) it replaces its original altogether (Christophel et al., 2015). Similarly, MEG recordings taken during mental rotation suggest that a gradual change occurs from the original representation into a rotated one (Trübutschek et al., 2019). By contrast, a recent study by Iamshchinina and colleagues (2021) showed that perceptual representations in primary visual cortex lasted throughout the mental rotation interval. It was nevertheless not the primary aim of these studies to track the possible simultaneous maintenance of perceived and transformed items in WM, so that the evidence is not clear-cut: In the studies by Christophel and colleagues (2015) and Trübutschek and colleagues (2019), the relationship between original and rotated items was constant across trials, which complicates any attempt to assess the presence of either item independently. For instance, between two representations rotated either 0 or 120 degrees, neural decoding of intermediate values cannot be definitely attributed to either representation. Conversely, in the study by Iamshchinina and colleagues (2021) there was only a single item being manipulated, so the original item may only have been found to persist as a consequence of presentation history, rather than it being part of WM proper.

A compounding, more general challenge to studying transformations in WM is the recent discovery that neural activity alone may not reflect the full breadth of WM operations. WM maintenance may not rely on unbroken chains of ongoing neural activity (LaRocque et al., 2013), as was previously thought (e.g., Curtis & D’Esposito, 2003; Kamiński et al., 2017). Rather, it may utilize activity-silent or quiescent brain states that are mediated by short-term changes in functional connectivity (Mongillo et al., 2008). Furthermore, it has been suggested that there may be different functional states in WM relating to passive maintenance and active, attentional updating (Olivers et al., 2011; Trübutschek et al., 2019), and both these states are presumably traversed when stored information is transformed. Importantly, although active attentional updating of WM is associated with easily measurable neural activity, this is not necessarily the case for passive maintenance (Kamiński & Rutishauser, 2020; but see also Muhle-Karbe, Myers, & Stokes, 2021; Stokes, Muhle-Karbe, & Myers, 2020). It is thus conceivable that traditional, activity-based measurement approaches miss part of the picture.

In our study, we sought to overcome these issues and decisively assess and compare WM states before and after mental transformation. First, although activity-quiescent states remain intrinsically difficult to measure non-invasively, it has recently been confirmed that functional connectivity can be illuminated by driving a standardized impulse signal through the network, as the response to that stimulation will partially reflect the momentary state of the network, independent from the focus of attention, and independent from its activity state (Buonomano & Maass, 2009; Stokes, 2015; Wolff et al., 2015; Wolff, et al., 2017; Rose et al., 2016). This is important, as EEG voltage decoding of working memory contents can drop to baseline within approximately a second (e.g., Fig. 2D in Wolff et al., 2017). Therefore, we implemented a perturb-and-measure approach (Wolff et al., 2015; 2017) by presenting task-irrelevant impulse stimuli during WM maintenance, before and after mental rotation of randomly oriented gratings. Second, we implemented a design in which original and transformed items were sufficiently independent from each other, so that they could be examined individually. Based on the research to date, two contrasting hypotheses were formulated: It may be that both the original and the transformed WM items are maintained and can be similarly decoded through impulse perturbation. Alternatively, one may hypothesize that WM only stores task-relevant information, so that once an item is transformed, only the resultant representation is kept, and the original item that is no longer relevant is rapidly discarded.

To preview the principal findings, before rotation, only the cued, relevant WM item could be successfully predicted from impulse-evoked EEG activity, unlike the uncued item, which seemed to have been rapidly purged from the WM system, replicating earlier findings (Wolff et al., 2017; 2020a; 2020b). Prior to the response probe, from the second impulse stimulus that followed the rotation instruction, the imagined rotation product could be decoded. Intriguingly, the original orientations could also still be decoded at that point in time. The continued presence of these obsolete originals in the WM network suggests that transformations in WM rely on relatively elaborate ‘double’ encoding, which was also associated with faster probe response times. Thus, the brain may prioritize representational precision and behavioural flexibility over storage capacity in spite of its scarcity.

## 2. Method

### 2.1 Participants

Thirty students of the University of Groningen (17 female, *M _Age_* = 21.3; *Range _Age_* = 18 – 31; all right-handed) volunteered to participate in the study in exchange for course credits or monetary reward (€8 per hour). The participants were selected from a larger group by means of a pre-screening procedure. Pre-screening consisted of a behavioural task, lasting approximately one hour, which was otherwise identical to the EEG session. The selection criteria were pre-determined and communicated to the participants; at least 70% task accuracy and a response time of less than 700 ms on average. The sample size was based on earlier studies with a similar design (e.g., Wolff et al., 2017). The study was conducted in accordance with the Declaration of Helsinki (2008), and was approved by the Ethical Committee of the Behavioural and Social Sciences Faculty of the University of Groningen (Study ID = 18029-SP). All participants provided written informed consent before taking part.

### 2.2 Apparatus and Stimuli

Participants were seated in a fully lit testing chamber at a viewing distance of approximately 60 cm from the screen, a 17” Samsung 797DF CRT monitor, set at a refresh rate of 100 Hz and a resolution of 1024 by 798 pixels. The stimuli were created and presented with the freely available Psychtoolbox 3 extension for Matlab (Brainard, 1997; Kleiner et al., 2007). A custom two-button response box connected via a USB interface was used to collect behavioural responses.

As shown in Fig. 1, the background remained grey throughout the experiment (RGB = 128, 128, 128), and a black fixation dot with a white outline (0.25° of visual angle) was present in the centre of the screen throughout the trials. The memory items were circles containing sine-wave gratings with 6 different orientations ranging from 15° to 165° with an interval of 30°. At the beginning of each trial, two orientation gratings were presented, each centred at 6.5° of visual angle (at 60 cm viewing distance) from the centre of the screen on the horizontal axis. All gratings were presented at 20% contrast, each again with a diameter of 6.5° of visual angle. The spatial frequency of the gratings was set to 0.65 cycles per degree, while their phase was randomized within and across trials. The cue stimuli were arrows of 0.5° of visual angle “>>”, pointing in the direction of the stimulus that was to be cued, presented in black Arial font 0.5° of visual angle above the centre of the screen. The direction of rotation was cued by an arrow of 1° of visual angle pointing clockwise or counter-clockwise (which was absent if no rotation was required), fitted to the middle of a 90° wide radian cut out from a circle with a diameter of 8° of visual angle. The angle of rotation was presented numerically (0, 30 or 60°) in bold black Arial font at a visual angle of 0.5° above the centre of the screen. The complete rotation instruction including the direction and the angle covered 2.75° of visual angle on the screen. The impulse stimulus consisted of three partially overlapping circles each with a diameter of 9.75°. The centre-to-centre distance of each circle was 6.5°. The probe stimulus was presented at the centre of the screen and was always identical to one of the 6 sine-wave gratings used as memory items.

**Figure 1.**
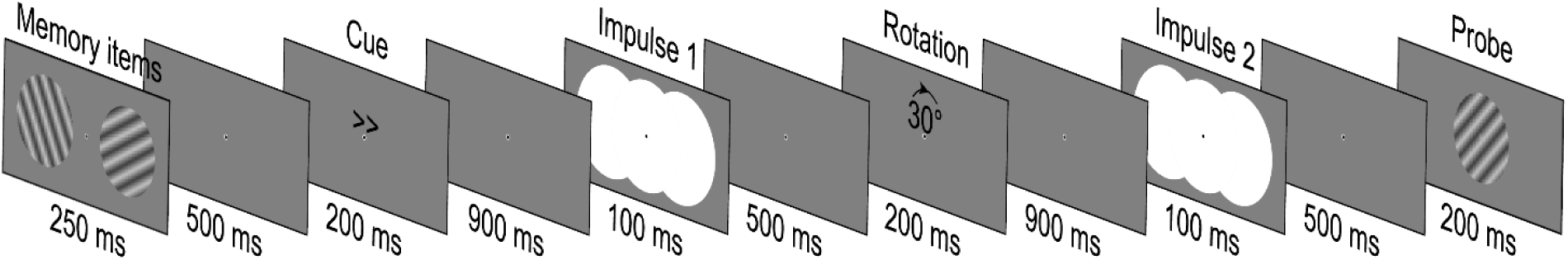
The design of a single experimental trial. Participants maintained one of two memory items in memory, indicated by a retro-cue. Subsequently, this item was rotated (0, 30, or 60°) and eventually compared to the probe stimulus. Impulse stimuli were task-irrelevant and served to elicit an EEG response by perturbing the representational network.

### 2.3 Procedure

Each participant first completed 48 practice trials that were identical to the experimental trials prior to EEG recording. During practice, all participants were instructed and trained to keep their gaze on the fixation dot at all times, and to blink only during or after response. Fast and accurate responses were encouraged. The whole experiment consisted of 1440 trials, spread across 4 consecutive sessions separated by breaks with a duration that was determined by the participants themselves. In each session, participants completed 24 blocks containing 15 trials each. Within these blocks, trials continued without interruption. On average, each participant took 4 hours to complete the task, including breaks.

Each trial began with the presentation of a fixation dot for 700 ms, followed by the presentation of the memory array, which consisted of two orientation gratings that appeared on both sides of the visual field for 250 ms. The orientations of the gratings were randomly selected without replacement from a uniform distribution, such that each of the six orientations was presented the same number of times throughout the experiment. Next, a blank screen with only the fixation dot was presented following stimulus offset for 500 ms. A retro-cue was then presented for 200 ms, indicating which of the two orientation gratings had to be retained in memory. The direction of the cue (left or right) was randomized, but evenly distributed across all conditions. After a delay of 900 ms the impulse signal was on display for 100 ms, and was followed by a blank delay interval of 500 ms. Then the rotation instruction was displayed for 200 ms, followed by another 900 ms delay. Like the cues, the rotation angles (0, 30 or 60°, clockwise and counter-clockwise) were randomized but evenly distributed across conditions. The second impulse was then presented for 100 ms, followed by a delay of 500 ms. Lastly, the probe was on display for 200 ms. Participants were asked to judge, as quickly as possible, whether the probe was the same as the relevant WM item, which was either the rotation product in rotation trials, or the original cued item on trials that did not require rotation (i.e., 0°). The probe was randomized but matched the relevant WM item in 50% of cases, while in the other 50% of cases, the probe was sampled randomly from the other possible orientations. The participants could report their answer by pressing one of the two buttons on the response box. After each response, feedback was given by a smiley that was presented for 200 ms, where a happy face indicated that the response was correct. The keys on the response box were counterbalanced across participants.

### 2.4 EEG Acquisition and Pre-processing

The EEG signal was recorded at a sample rate of 1000 Hz from 62 Ag/AgCl sintered electrodes deployed with a 10-20 international layout. The average of all electrodes was used as the reference during recording, with a ground electrode placed on the sternum. The data were recorded with BrainVision Recorder software, and a TMSI Refa 8-64/72 amplifier. Eye movements were tracked via bipolar electrooculography with vertical electrodes above and below the left eye and two horizontal electrodes on ipsilateral sides of both eyes. The impedance at all electrodes was kept below 10 kΩ.

Offline, the data were re-referenced to the mastoids, downsampled to 500 Hz and bandpass filtered (0.1 Hz high-pass and 40 Hz low-pass) using EEGLAB (Delorme & Makeig, 2004). The data were epoched relative to Impulse 1 and Impulse 2 (-150 ms to 600 ms). Epochs with excessive variance, voltage drifts, or muscle or eye movement artefacts were identified visually and removed from all subsequent analyses. In total, 11.3% of epochs were rejected.

Finally, the data were reformatted to best capture the time-locked neural dynamics evoked by the impulse stimuli. The approach was the same as established previously (Wolff et al., 2020a, 2020b), and the experiment was designed to take advantage of it. For the full *spatiotemporal* analyses, the voltage traces from the 17 posterior channels of interest (P7, P5, P3, P1, Pz, P2, P4, P6, P8, PO7, PO3, POz, PO4, PO8, O1, Oz and O2) were extracted relative to the impulse onsets from 100 to 400 ms. Then, the mean voltage within each time window was removed for each trial and channel separately, which removes drift (as in conventional baselining) and isolates the dynamic, stimulus-evoked neural response. The voltage traces were then down-sampled to 100 Hz, and combined with the channel dimension, resulting in 510 data features (30 time-points x 17 channels) per trial for each impulse. The primary advantage of this method is that it does not require a pre-stimulus baseline that is either not neutral (in the case of taking the baseline right before the impulse), or temporally too far away to be effective due to signal drift (in the case of a pre-stimulus baseline). As the method ensures that only data from the time-window in question is used, the relative baselines used here for Impulse 1 and 2 are equally effective, compared to a pre-trial baseline that could be ‘better’ for Impulse 1 (because it is temporally closer) than for Impulse 2, prohibiting a direct comparison between them, since reported effects could be ‘worse’ at Impulse 2 due to more unremoved drift compared to Impulse 1. While a higher higher-pass filter might alternatively be used to effectively remove drift and negate the need for a baseline altogether, this could in turn introduce temporal distortions in the data (van Driel, Olivers, & Fahrenfort, 2021), similar to a non-neutral baseline. A further advantage of the present method is that it may improve sensitivity by combining both temporal and spatial dimensions of the dynamic signal.

For the *time-course* analyses of neural dynamics, the same posterior channels as above were included. Here we used a 100 ms sliding window from -50 ms to 550 ms relative to impulse onsets. Similar as above, the data were down sampled to 100 Hz within each time-window, and the mean voltage was removed. The resulting 10 time-points were then combined with the channels, resulting in 170 data features. The analyses (described below) were done on each time-point that the time-window was centred on, separately. The time-courses were smoothed with a gaussian smoothing kernel (*SD* = 16 ms).

For the *searchlight* analyses, the whole spatiotemporal window was used, but the analyses were repeated iteratively for each electrode and its closest two neighbours (thus 30 time-points x 3 channels), across all 62 electrodes.

### 2.5 EEG Analyses

#### 2.5.1 Linear discriminant contrast

We used cross-validated representational similarity analysis (RSA) with Mahalanobis distance (Nili et al., 2014), also termed linear discriminant contrast (LDC; Walther et al., 2016), to investigate the contributions of different task-related models to the neural codes evoked by Impulse 1 (before rotation instructions) and Impulse 2 (after rotation instructions). We used an 8-fold cross-validation approach to compute the squared Mahalanobis distance (MD^2^) between condition pairs of all condition combinations of interest at Impulse 1 and Impulse 2, using the following formula:

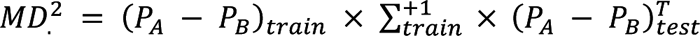

The data was randomly split into 8 folds using stratified sampling that ensures a roughly equal number of trials in each fold. The *test* data consisted of a single left out fold and the *train* data consisted of the remaining 7 folds, which was also used to estimate the noise covariance matrix (∑). The number of trials of each condition was equalised through random subsampling within the test fold and train folds. P_A_ and P_B_ were the trial-averaged patterns of conditions A and B of the subsampled trials of the *test* and *train* data. The noise covariance matrix was estimated from the subsampled *train* data by subtracting the trial-averaged activity patterns of each condition from all trials of the corresponding condition. The covariance matrix was calculated using a shrinkage estimator (Ledoit & Wolf, 2004) on the resulting condition-mean centred trial by feature matrix. The pseudoinverse of the covariance matrix (∑^+1^) was then used to compute the MD^2^ between all possible condition differences resulting in an n-conditions by n-conditions representational dissimilarity matrix (RDM). This procedure was repeated for all test and train fold combinations. The 8-fold partitioning and the subsampling within test and train data was randomised. To ensure stable results, the above procedure was repeated a total of 100 times for each subject, with random folds and random subsampling each time. The resulting data RDMs were averaged across repetitions and folds, resulting in a single RDM per subject and impulse. These RDMs were z-scored and then regressed against specific task models of interest.

The conditions that were considered and the task models were different for each impulse. At Impulse 1 we considered the cued item (6 orientations), the uncued item (6 orientations), and the cued location (left or right). The pairwise MD^2^s between all 72 unique condition combinations resulted in a 72 × 72 data RDM for each subject. The task models we tested at Impulse 1 comprised the cued location model (left or right), parametric coding models for the cued and uncued memory item (absolute circular distance between cued items, and between uncued items), as well as a parametric coding model of the generalization between cued and uncued items (absolute circular distance between cued and uncued). The parametric coding models thus assumed a circular relationship between absolute circular distances and pattern similarities. All models (except the cued location model) were subdivided and tested separately within the same and across different cueing conditions to test for (cued) location specificity. For example, when testing only for the same cued location version of the cued item model, the cells comprising across-location generalization (which were in our case in the upper left and lower right of the design matrix) were down-weighted to 0 by replacing them with the average of the design matrix. All task model RDMs were z-scored and regressed simultaneously against the z-scored data RDM of each subject using multiple regression.

At Impulse 2 we considered the cued item (6 orientations), the rotation instructions (5 conditions), and the cued location (left or right), which resulted in a 60 × 60 RDM for each subject. We first tested the cued location and the rotation instructions task models simultaneously using multiple regression. The rotation instructions models consisted of 5 separate models (one for each condition), each testing same vs. different instructions. For example, for rotation condition -60 degrees, the model consisted of a matrix that assumed higher similarity/lower Mahalanobis distance for all condition-combination cells that included the rotation condition -60 degrees, which were filled with -1s, whereas all other cells were filled with 1s. The same logic applied to all other rotation condition models. We did thus not make any assumptions about the relationship between the rotation conditions. The resulting model fits/beta values of the rotation models were averaged for plotting and statistical analyses. We furthermore tested parametric coding models of the cued item, the rotated item, and the generalization between them using a similar logic as for the parametric coding models used for Impulse 1 described above. However, since the cued and the rotated item models are statistically related, the models were fit on the residual data RDMs of the other. This meant that the cued item model was fit to the residual variance of the data RDM that was not accounted for by the rotated item model data, and vice versa. The rotated item model was marginally related to the rotation instructions model, so it too was regressed out before fitting the rotated item model. The cued-rotated generalization model was fit to the residual variance of the data RDM that was not accounted for by either the cued item model or the rotated item model. As in the case of Impulse 1, all models (except for the cued location model) were subdivided and tested separately for within and across cued location conditions. All model and data RDMs, as well as the residual variance RDMs, were z-scored before regression.

Note that we ran simulations (see “**Simulations of neural patterns**”) that included the same dependence between cued and rotated item when including all rotation conditions, which confirmed that fitting the models on the residuals adequately removed any residual effects. For example, when only the rotated item is modelled, fitting the cued model on the residual showed no effect for the cued item (see Fig. 4a). We believe that the used RSA approach was thus appropriate, enabling us to test the contribution of different effects but related models, without the need remove data. Nonetheless, we repeated the RSA at Impulse 2 after excluding all no-rotation trials, thus removing the dependency between cued and rotated items. This resulted in a 48 by 48 RDM (2 locations, 4 rotation conditions, 6 cued item orientations). The models were the same as described above (minus one rotation condition model due to the exclusion). However, the models of either the cued or the rotated item were not fit on the residual of the other since they were not statistically related to one another anymore, rendering this step unnecessary.

Finally, as an additional control, we also applied an RSA with the rotated item models and the cued item models to the full data at Impulse 2, without trial removal. Crucially however, the model fit related to one model was estimated independently before it was regressed out. We first randomly split the data into two independent halves (data_1 and data_2), before computing the MD^2^s between all trial condition combinations within each half independently using 4-fold cross-validation, resulting in two 60 by 60 RDMs (RDM_1 and RDM_2) for each subject. The cued model fit estimated from RDM_1 was then regressed out from RDM_2, before fitting the cued and the rotated item models to the now residual RDM_2. The same was done for the rotated item model fit, which was estimated from RDM_1, regressed out from RDM_2, before again applying both models to the residual RDM_2. The 1000 beta values for each model fit on each residual RDM per subject were then averaged for subsequent statistical significance testing. Given the presence of both items in the data, if the cued item model contribution estimated from data_1 selectively removed only the cued item effect from data_2, without affecting the rotated item effect, then the average cued item model fit to the residual RDM_2 should, on average, be 0, while the rotated item fit should still be positive (and vice versa when regressing out the rotated item). This procedure was repeated 1000 times, with random data splits in each iteration.

### 2.5.2 Correlation of trial-wise decoding strength

We are not aware of a trial-wise RSA approach that can test the contribution of correlated models. Thus, in order to test if and to what extent the trial-wise strength in neural activation patterns related to the cued item correlated with the corresponding neural activation patterns of the rotated item, we used the same trial-wise decoding approach as in Wolff et al. (2020b) at Impulse 2. For this analysis we excluded all 0 rotation trials to ensure independence between cued and rotated items (note that the same was done for the simulations). This entailed using an 8-fold cross-validation decoding approach using Mahalanobis distance. The data was randomly split into 8 folds using stratified sampling (ensuring a roughly equal number of trials for each orientation in each fold). The covariance matrix (with shrinkage estimator; (Ledoit & Wolf, 2004)) was computed using the 7 folds of the train data. The number of trials of each orientation within train data was then equalised via random subsampling. The averaged patterns of each orientation of the train data were convolved with a half cosine basis set raised to the 5^th^ power to pool information across similar orientations (Myers et al., 2015). The Mahalanobis distance between each test trial and averaged orientation patterns of the train data were then computed. This procedure was repeated for all test and train fold combinations. The resulting 6 distances for each trial were summarized into a single “decoding strength” value by computing the cosine vector mean of the absolute circular distance between test trial’s orientation and averaged orientation patterns of the train trials. To get reliable estimates, the above procedure was repeated 100 times (with random folds and subsamples), resulting in 100 decoding strength values for each trial and subject, which were subsequently averaged. The cued and the rotated items were decoded separately at Impulse 2. The Pearson correlation between the trial-wise decoding strengths of the cued and the rotated item was computed for each subject separately, Fisher’s z-transformed, and statistically tested against 0 (see “**Statistical significance testing**”).

#### 2.5.3 Decoding generalization before and after mental transformation

We were interested if the neural pattern related to the cued item before mental rotation (at Impulse 1) generalized to the neural pattern after mental rotation (at Impulse 2). To test this, we trained a classifier using the same approach as described in the previous section on the neural pattern in all the trials of the cued item at Impulse 1 and tested it only on the rotation trials at Impulse 2 (i.e., excluding the trials in which subjects were instructed not to rotate the item), using 8-fold cross-validation, random subsampling within test and train folds, and 100 repetitions. Given the cue-specific coding scheme observed at Impulse 1, the classifier was trained and tested within each cued location condition separately and averaged afterwards. The test trials at Impulse 2 were either labelled with the cued item, to test temporal generalization of the cued item over time (Wolff et al., 2020b), and after mental rotation, or labelled with the rotated item, to test whether the same neural pattern that codes for the cued item was re-used to code for the rotated item.

#### 2.5.4 Relationship between the neural and behavioural data

We were interested if the quality of WM content predicted behavioural performance (probe response times and accuracy) on a trial-by-trial basis within subjects. We used the trial-wise decoding strengths of the cued/original and the rotated item at Impulse 2 (excluding the “0 rotation” trials) and tested if they predicted trial-wise fluctuations in accuracy and response times. First, we regressed the decoding strengths against each other to obtain the residuals of each, to ensure that the decoding strengths of the cued/original and the rotated item were uncorrelated and explained unique aspects of the behavioural measure in question. We then used the residuals of the cued/original and the rotated item decoding strengths as regressors to predict behavioural accuracy (logistic regression) and log-transformed response times (linear regression) within each subject. The resulting regression weights were then tested for significance in the expected direction of facilitation.

### 2.6 Simulations of neural patterns

We simulated several plausible effects to compare the pattern of results of simulated data with the results of the actual data, and to ensure that our analyses are adequate for our experimental design. Activity patterns were simulated by randomly drawing 20 values from the standard normal distribution two times. To simulate a parametric pattern for the circular memory items, one of the two patterns was convolved with the sine of the memory item in question, and the other with the cosine of the same memory item of that trial, before adding both signals together. Same coding schemes for each item (original and rotated) were simulated by using the same random pattern for the sine, and the same random pattern for the cosine, of the orientation of each item. Unique coding schemes were simulated by using randomly different random patterns for the sine and cosine of the orientation of each item.

Trial-wise noise was added to the signal by randomly drawing 20 values from the standard normal distribution and then multiplying it with a random value drawn from a normal distribution with mu = 10 and SD = 3. This was done separately for each trial to simulate trial-wise fluctuations in neural noise levels. The trial-wise noise patterns were added to the signal patterns comprising the overall simulated neural signal. We used the same number of trials and proportion of conditions as the experiment, including 0 rotation trials.

We simulated and analysed the following scenarios that we thought could be expected in this task:

A. Rotated only: A new activity pattern for the rotated item is present in the signal, while the code of the original item completely disappears.
B. Partial rotation: An activity pattern is present in every trial that represents an item that is only partially rotated (half-way). Neither the original, nor the fully rotated item are represented in the signal. This simulates the possibility that subjects did not fully mentally rotate the item, and/or that its orientation is attracted to the orientation of the original item.
C. Same coding schemes: The original and the rotated item use the same signal pattern and both are present in the data.

i. The patterns of both items are simultaneously present in every trial.
ii. The pattern of only one of the items is present in a given trial, simulating the possibility that subjects may have only sometimes followed instructions and mentally rotated the item.
D. Unique coding schemes: The original and the rotated items use unique and independent coding schemes and are both present in the data.

i. Both patterns present in every trial.
ii. Only the pattern of one of the items is present in a given trial.

For simplicity we did not consider cue-specific effects in the simulations (i.e., whether or not the coding schemes generalize across cued-locations or not). We ran each scenario 100 times, with randomised signal and noise patterns each time (simulating 100 subjects). The simulated data was subsequently analysed in the same way as the real data of Impulse 2 (LDC and correlation of trial-wise decoding strengths).

We re-ran the above simulations with explicit rotation condition signals added to them. We wanted to ensure that rotation signals do not alter the effects related to the original and rotated item. The rotation condition signal consisted of 6 (one for each rotation condition) unique random patterns (20 values drawn from a normal distribution), one of which was added to the simulation signal in each trial, depending on the rotation condition of that trial.

### 2.7 Statistical significance testing

We used non-parametric tests to assess statistical significance in all cases, and all statistical tests were two-sided (unless explicitly stated otherwise). We tested for statistical significance of the neural analyses results by randomly shuffling the conditions in question 1,000 times and using the resulting null-distribution to conduct a *t*-test. In the case of the LDC and trial-wise decoding analyses this meant that the analyses were re-run with randomised condition labels resulting in 1,000 “null” model fits/decoding values per subject. These were transformed into null distributions of *t* values by computing the t-value across subjects for each of the 1,000 values, which was then used for a *t* test against 0 of the actual model fit/decoding value. For the time-course analyses, a cluster-based permutation test (1,000 permutations) was used to correct for multiple comparisons over time with a cluster-forming threshold of *p* < 0.05.

To test for statistical significance of the correlation between trial-wise decoding values of the cued and the rotated item, the trial-wise decoding values were randomly shuffled 10,000 times, each time obtaining the Fisher’s z-transformed correlation value. These were transformed into a t-value distribution which was then used to perform a t-test.

The relationship between neural and behavioural data was tested for statistical significance by shuffling the trials 10.000 times and repeating the regression analyses each time. The resulting null-distribution of the beta values was used to perform a t-test.

We also computed Bayes Factors (BF) to complement p-values. We used the Bayesian implementation of the non-parametric Wilcoxon signed-rank test with 10.000 samples and the Cauchy prior with the default scale of 0.707, as implemented in JASP (JASP Team, 2018).

### 2.8 Data and code availability

All data and MATLAB code used to generate the results and figures of this manuscript are publicly available at https://osf.io/3hdpc and at https://github.com/mijowolff/veridical-and-transformed-representations-in-wm, respectively.

## 3. Results

### 3.1 Behavioural Results

Behavioural measures consisted of response accuracy (% correct), reflecting the comparison of the probe and the memory item, and reaction time (RT), reflecting the time from probe onset until the button press. Mean accuracies and median RTs were analysed as a function of rotation condition (-60, -30, 0, 30, 60). Behavioural analyses were conducted in the freely available JASP program (JASP Team, 2018).

The results were typical for rotation tasks (e.g., Wexler et al., 1998; Searle & Hamm, 2017), and are shown in Fig. 2. The accuracy in the task was highest when no rotation took place and declined as the rotation magnitude increased, regardless of direction. Repeated measures ANOVA confirmed that accuracy in at least one condition differed from the others, *F*(4, 116) = 61.454, *p* < 0.001, *η^2^* = 0.679, *BF_10_* > 1000. Post hoc paired *t*-tests revealed that performance differed between all rotation conditions, unless rotation magnitude was identical (*t* _-60,-30_ (29) = -5.127, *p* < 0.001, *BF_10_* = 588.67; *t* _-60, 0_ (29) = -12.613, *p* < 0.001, *BF_10_* > 1000; *t* _-60, 30_ (29) = -5.589, *p* < 0.001, *BF_10_* > 1000; *t* -_60, 60_ (29) = 1,444, *p* = 1, *BF_10_* = 0.476; *t* _-30,0_ (29) = -7.487, *p* < 0.001, *BF_10_* > 1000; *t* _-30, 30_ (29) = -0.463, *p* = 1, *BF_10_* = 0.217; *t* _-30, 60_ (29) = 6.570, *p* < 0.001, *BF_10_* > 1000; *t* _0, 30_ (29) = 7.024, *p* < 0.001, *BF_10_* > 1000; *t* _0, 60_ (29) = 14.057, *p* < 0.001, *BF_10_* > 1000; *t* _30, 60_ (29) = 7.033, *p* < 0.001, *BF_10_* > 1000; Bonferroni-corrected; Fig. 2A). The distribution of accuracies across rotation conditions clearly showed that greater rotation magnitudes negatively influenced recall accuracy, though accuracy remained well above chance in all cases.

**Figure 2.**
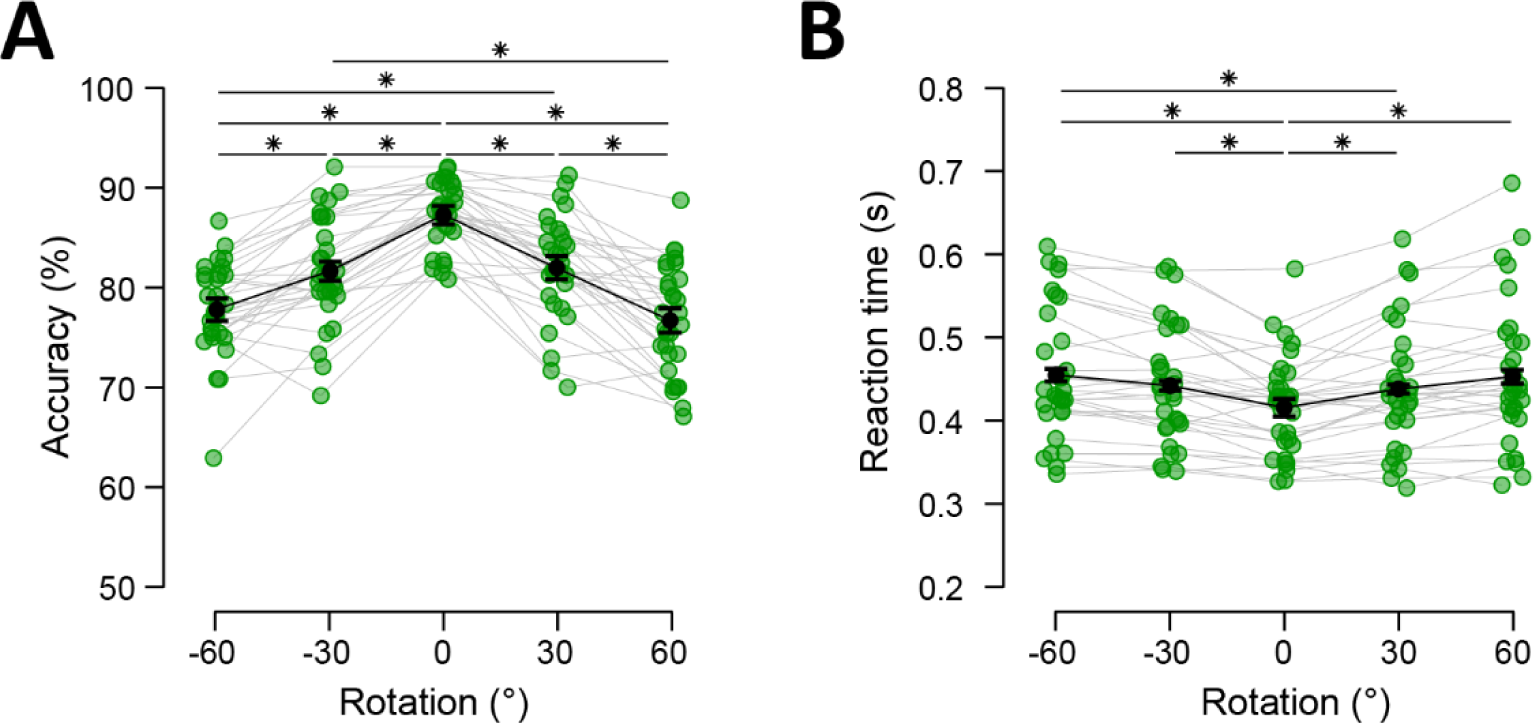
Behavioural performance as a function of rotation. **(A)** Mean accuracy in percent correct. **(B)** Reaction times (means of medians) in seconds. Green dots represent individual data points, black dots reflect averages, and error bars represent within-subject 95% confidence intervals (Morey, 2008). Significant pairwise differences are indicated with asterisks (* *p* < 0.05, Bonferroni-corrected).

Rotation magnitude similarly influenced RT, *F*(2.601, 75.429) = 16.989, *p* < 0.001, *η^2^* = 0.369, *BF_10_* > 1000 (with Greenhouse-Geisser correction), with an increase in RT as rotation magnitude increased (*t* _-60,-30_ (29) = -2.376, *p* = 0.191, *BF_10_* = 10.993; *t* _-60, 0_ (29) = 7.282, *p* < 0.001, *BF_10_* > 1000; *t* _-60, 30_ (29) = 3.119, *p* = 0.023, *BF_10_* = 26.003; *t* -_60, 60_ (29) = 0.367, *p* = 1, *BF_10_* = 0.212; *t* _-30, 0_ (29) = 4.905, *p* < 0.001, *BF_10_* = 438.915; *t* _-30, 30_ (29) = 0.743, *p* = 1, *BF_10_* = 0.353; *t* _-30, 60_ (29) = -2.009, *p* = 0.468, *BF_10_* = 1.013; *t* _0, 30_ (29) = -4.163, *p* < 0.001, *BF_10_* = 80.012; *t* _0, 60_ (29) = -6.915, *p* < 0.001, *BF_10_* > 1000; *t* _30, 60_ (29) = -2.752, *p* = 0.069, *BF_10_* = 14.801; Bonferroni-corrected; Fig. 2B). This clear decrease in performance as a function of rotation magnitude strongly suggests that items were indeed mentally rotated and not replaced from a fixed stimulus set from long-term memory.

### 3.2 EEG results

#### 3.2.1 LDC at Impulse 1; before transformation/after cue

The RDM of Impulse 1, depicting the MD^2^ between all cued, uncued, and cued location combinations, is shown in Fig. 3A. The cued location model showed a significant effect (spatiotemporal: *p* < 0.01, BF_10_ > 1000; time-course: *p* < 0.01, -55 ms to 550 ms, cluster-corrected, Fig. 3B), likely driven by a shift in spatial attention toward the cued item in WM (e.g., Wolff et al., 2017). The cued item coding model was statistically significant within cued location (spatiotemporal: *p* = 0.02, BF_10_ = 11.117; time-course: *p* < 0.01, 92 ms to 300 ms, cluster-corrected), cluster-corrected, but not across (spatiotemporal: *p* = 0.712, BF_10_ = 0.219), and the difference between within and across cued location conditions of the cued item coding model was statistically significant (spatiotemporal: *p* = 0.042; time-course: *p* < 0.01, 140 ms to 292 ms, cluster-corrected; Fig. 3C, left), though the Bayesian evidence for or against an effect was ambiguous (BF_10_ = 1.671). The uncued item coding models showed no significant effects and Bayesian evidence against effects (spatiotemporal, within cued location: *p* = 0.354, BF_10_ = 0.276; across cued location: *p* = 0.426, BF_10_ = 0.242; difference: *p* = 0.242, BF_10_ = 0.394; Fig. 3C, middle). These results replicate previous findings in EEG of cue-specific neural coding of the cued WM item when it was laterally presented, and no detectable trace of the uncued item (Barbosa et al., 2021; Wolff et al., 2017; 2020a; 2020b). None of the generalization models between the cued and uncued item reached the statistical significance threshold (spatiotemporal, within cued location: *p* = 0.536, BF_10_ = 0.238; across cued location: *p* = 0.058, BF_10_ = 1.349; difference: *p* = 0.368, BF_10_ = 0.257; Fig. 3C, right).

**Figure 3.**
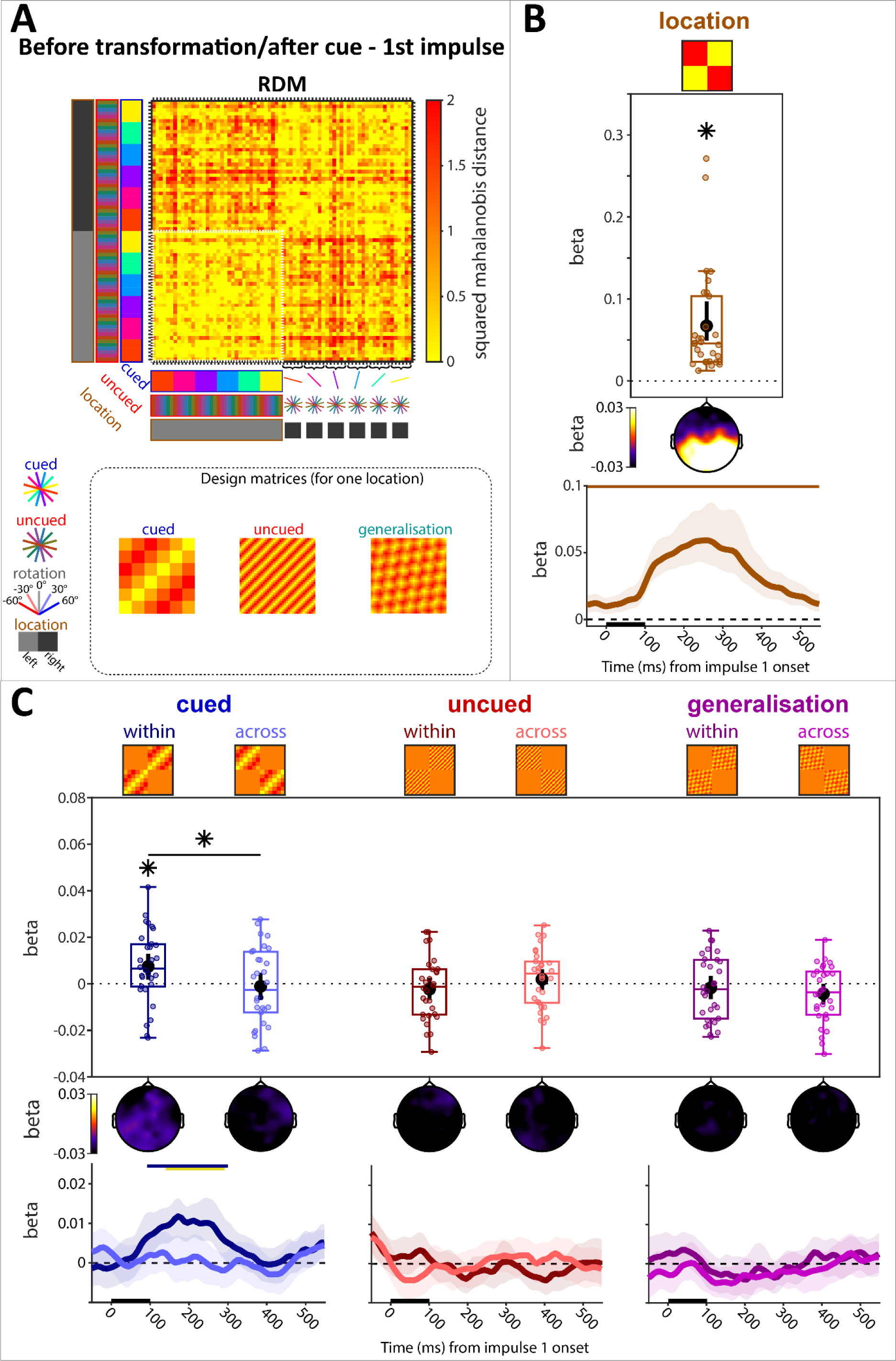
LDC before transformation & after the retro-cue at Impulse 1. **(A)** Average RDM and design matrices. Condition combinations are color-coded. All condition combinations for the right cue condition are explicitly illustrated on the y-axis. The remaining cells follow the same convention. The inset shows the design matrix for each model (except cued location) for one cued location (the size of the white outline in the RDM **(B)** Model fit (beta) of cued location condition. Top: Spatiotemporal; Middle: Searchlight; Bottom: Time-resolved. **(C)** Model fits of WM items (cued, uncued) and their generalization. Models are separated into within-cued location condition (within) and across-cued location condition (across). Same convention as (B). Error bars (spatiotemporal) and error shadings (time-resolved) are 95% C.I. Statistically significant (*p* < 0.05) model fits and differences of model fits between within and across locations are marked with (*). Statistically significant time-course clusters (*p* < 0.05, cluster-corrected) are marked with coloured bars at the top (corresponding colour for significant model fits, yellow for differences between within and across locations).

We used a relative, within time-window baseline (see “**EEG Acquisition and Pre-processing**”), as used in previous works (Wolff et al., 2020a, 2020b). The pattern of results stayed the same when using a pre-trial baseline instead, though the generalization model between cued and uncued item across cued location reached the significance threshold when using a pre-trial baseline (Suppl. Fig. 1A).

#### 3.2.2 Simulations of predicted neural effects after mental transformation in WM

We simulated plausible changes in the neural coding schemes of WM content after mental transformation. The analyses for the simulated data and the data at Impulse 2 were the same as well as using largely the same models, though for simplicity, we did not simulate or model spatial specificity, that is, differences in coding schemes when the original item was presented on the left or the right side. The pattern of results for each scenario are shown in Fig. 4. We simulated the presence of only the code for the rotated item (Fig. 4A), a single code for an orientation that is halfway between the original and the fully rotated item (i.e., partial rotation, Fig. 4B), as well as the presence of both the original and the fully rotated items simultaneously (Fig. 4C & D). Either the coding schemes for both were the same (i.e., using the same random patterns to represent the orientations of each item, Fig. 4C), or different (i.e., using randomly different random patterns to represent the orientations of each item, Fig. 4D). Note that almost every considered scenario resulted in a qualitatively unique pattern of results. The exception is the partial rotation of the original item (Fig. 4B) and the simultaneous representation of both the original and the rotated item in every trial, both using the same coding schemes (Fig. 4Ci), which cannot be distinguished with the analyses we employ. This means that if subjects systematically only partially rotated the item, resulting in an attraction of the orientation of the rotated item to the orientation of the original item, the results may falsely imply that both items were simultaneously present in every trial. However, as seen in the next section below, the pattern of results of the actual data at Impulse 2 after transformation best resembled the scenario depicted in Fig. 4Di: the simultaneous maintenance of both items in every trial (original and rotated), with unique coding schemes.

**Figure 4.**
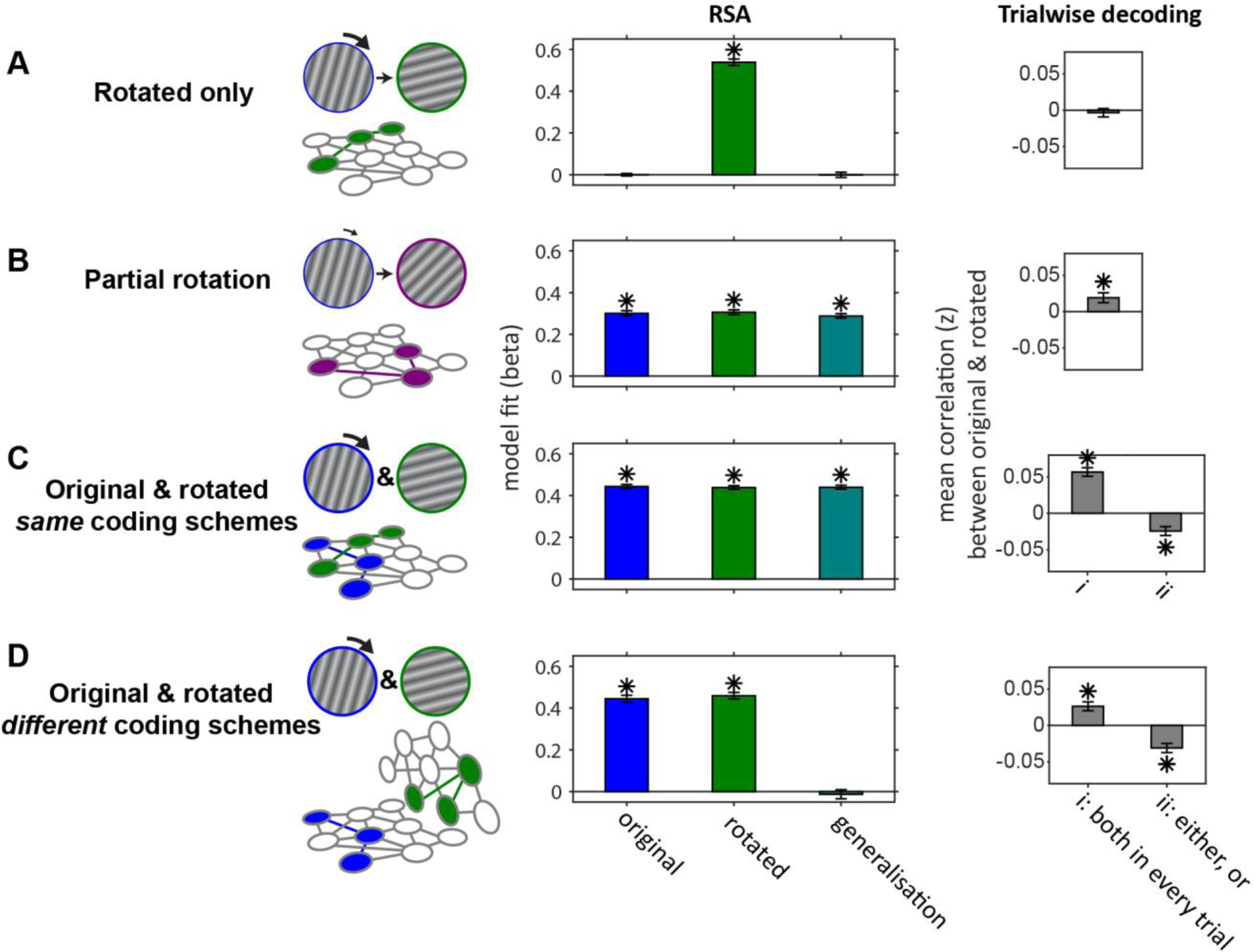
Simulation results (*N* = 100) of plausible maintenance scenarios of the original cued item and the rotated item after mental transformation. The left panel shows the schematics of each scenario, the middle panel the model fits of the original item, the rotated item, and the generalisation between them, and the right panel the mean correlation (Fisher’s z) between the trial-wise decoding strengths of the original and the rotated item. **(A)** Rotated only: Only the fully rotated item is present in the WM network, and the original item is removed completely. **(B)** Partial rotation: The original item is only halfway rotated in each trial and the original item is no longer present. Note that the analyses would incorrectly imply that both the original and the fully rotated items are simultaneously present in the network using the same coding schemes (i.e., generalisation between them) **(C)** Original & rotated, *same* coding schemes: The original and the (fully) rotated items are both coded using the same coding schemes. i) both are present in every trial (note the same pattern of results as B). ii) either one, or the other is represented in a given trial. **(D)** Original & rotated, *different* coding schemes: The original and the rotated item are represented in unique coding schemes, i) simultaneously on every trial, or ii) only one of them is randomly represented on every trial). Error bars are 95% C.I. Asterisks denote statistically significant (*p* < 0.05) results of the simulated data.

None of the simulations reported in Fig. 4 included an explicit signal that codes for the rotation conditions, and accordingly, none of the model fits were significant (Suppl. Fig. 2, left column). Including an explicit rotation condition signal in the simulations, unsurprisingly, resulted in significant model fits for the rotation condition model, but did not alter the pattern of results for the remaining model fits reported in Fig. 4 (Suppl. Fig. 2, right column).

#### 3.2.3 LDC at Impulse 2; after transformation

The RDM of Impulse 2, depicting the MD^2^ between all cued item, rotation instructions, and cued location condition combinations, is shown in Fig. 5A. The cued location model was significant (spatiotemporal: *p* < 0.01, BF_10_ > 1000; time-course: *p* < 0.01, 68 ms to 452 ms, cluster-corrected), as were both rotation instruction coding models (spatiotemporal, within cued location: *p* < 0.01, BF_10_ > 1000; time-course: -55 ms to 20 ms, 52 ms to 550 ms, *p* < 0.01, cluster-corrected; spatiotemporal, across cued location: *p* < 0.01, BF_10_ > 1000; time-course: 36 ms to 550 ms, *p* < 0.01, cluster-corrected), with no difference between them (spatiotemporal: *p* = 0.478, BF_10_ = 0.326; Fig. 5B). Even though the original orientation of the cued item was not behaviourally relevant at Impulse 2 anymore, both cued item coding models were statistically significant for both within cued location (spatiotemporal: *p* < 0.01, BF_10_ = 11.126; time-course: 236 ms to 428 ms, *p* < 0.01, cluster-corrected) and across cued location (spatiotemporal: *p* = 0.044, BF_10_ = 1.454; time-course: 108 ms to 148 ms, *p* = 0.012, 268 ms to 308 ms, *p* = 0.044, cluster-corrected Fig. 5C, left), though Bayesian evidence was ambiguous for the latter. This suggests that the neural code of the cued item was less spatially specific at Impulse 2 after transformation, in contrast to the code at Impulse 1 before transformation. However, while there was no difference between them in the spatiotemporal analysis (*p* = 0.448, BF_10_ = 0.248), the time-course analysis showed a significant difference between within- and across-models of the cued item (332 ms to 380 ms, *p* < 0.01, cluster-corrected), suggesting some spatial specificity, nonetheless.

**Figure 5.**
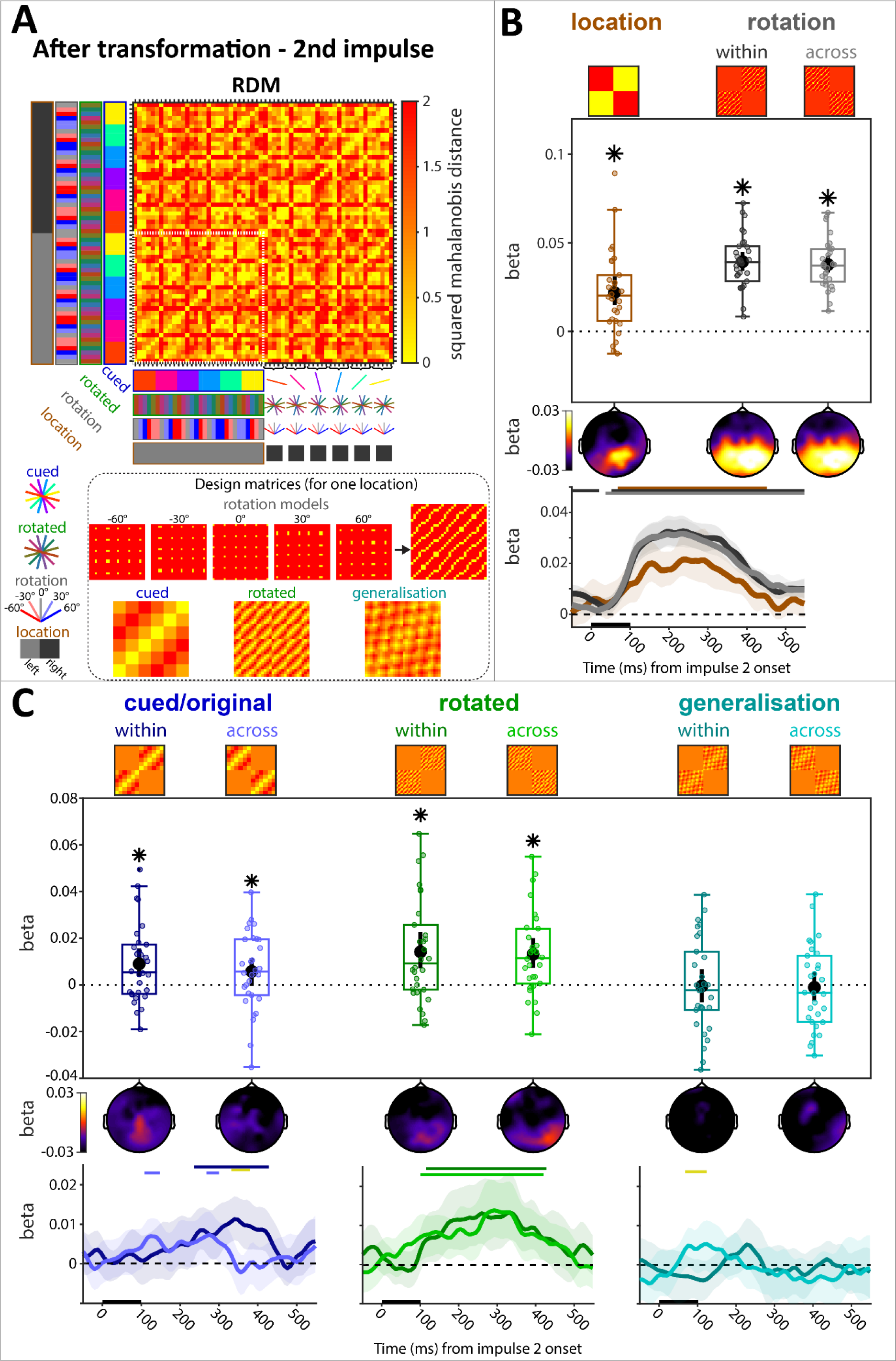
LDC after transformation at Impulse 2. **(A)** Average RDM and design matrices. Condition combinations are color-coded. All condition combinations for the right cue condition are explicitly illustrated on the y-axis. The remaining cells follow the same convention. The inset shows the design matrix for each model (except cued location) for one cued location (the size of the white outline in the RDM). **(B)** Model fit (beta) of cued location condition and rotation condition (separately for within cued location and across cued location). Top: Spatiotemporal; Middle: Searchlight; Bottom: Time-course. **(C)** Model fits of the original cued item, the rotated item, and their generalization. Models are separated into within cued location condition (within) and across cued location condition (across). Same convention as (B). Error bars (spatiotemporal) and error shadings (time-course) are 95% C.I. Statistically significant (*p* < 0.05) model fits and differences of model fits between within and across locations are marked with asterisks (*) for spatiotemporal. Statistically significant clusters (*p* < 0.05, cluster-corrected) are marked with coloured bars at the top (corresponding colour for significant model fits, yellow for differences between within and across locations) for the time-course.

The rotated item coding models were both statistically significant (within cued location, spatiotemporal: *p* < 0.01, BF_10_ = 53.742; time-course: 116 ms to 428 ms, *p* < 0.01, cluster-corrected; across cued location, spatiotemporal: *p* < 0.01, BF_10_ = 251.767; time-course: 100 ms to 420 ms, cluster-corrected), and were not different from each other (spatiotemporal: *p* = 0.68, BF_10_ = 0.196; Fig. 5C, middle). The generalization coding models between the cued and the rotated item were not significant with Bayesian evidence for no effect (spatiotemporal: within cued location: *p* = 0.892, BF_10_ = 0.198; across cued location: *p* = 0.704, BF_10_ = 0.210; difference: *p =* 0.912, BF_10_ = 0.213; Fig. 5C, right), though the time-course analysis revealed a significant cluster in the difference (time-course: 68 ms 124 ms, *p* < 0.01, cluster-corrected).

We explicitly tested if there was a significant difference between the within-item models (cued and rotated) and the generalization models, which would provide evidence that the cued and the rotated items are coded in significantly different coding schemes from one another. To quantify this “cost of generalization” (cf. Wolff et al. 2020b), we compared the average beta values of the within-item models with the average of the generalization models. The difference between within-item models and the generalization models was significant (spatiotemporal: *p* < 0.01, BF10 = 14.157).

The pattern of results of the spatiotemporal signal when applying a pre-trial baseline instead of a relative, within time-window baseline, was qualitatively the same (Suppl. Fig. 1B).

We also explored whether decoding was possible from stable delay activity that is minimally influenced by the presentation of the impulse, by using the averaged voltage traces right before Impulse 2 (-100 to 0ms), and after (500 to 600ms), when the impulse-evoked signal has presumably largely subsided. Cued items could be decoded, but there was no evidence for the rotated item from neural activity that was not evoked by the impulse (Suppl. Fig. 3), suggesting that the impulse may be critical to ‘illuminate’ WM content. However, this result should be considered with caution, as the present study was not designed to directly measure the impact of the impulse on WM decoding.

We furthermore checked whether the pattern of results of the spatiotemporal signal remained the same, when excluding all no-rotation trials. The results remained largely the same (Suppl. Fig. 4), though the across-location model of the cued item failed to reach significance.

Finally, when keeping the 0 rotation trials in the data, we found that when the effect of one model (cued or rotated item) was estimated from independent data before regressing it out from the test data, only the fit of the model that was not removed was statistically significant (Suppl. Fig. 5).

Overall, these results provide evidence for the presence of both the cued, and the mentally rotated items in the WM network, which are both coded using distinct coding schemes that do not cross-generalize, in line with the simulation results of scenario 4D above.

#### 3.2.4 Correlation between trial-wise decoding strengths of the original and the rotated item after transformation

The results presented above are based on trial-averaged data and do not rule out the possibility that although evidence for both the cued/original and the rotated item was found, subjects only maintained one of the two in individual trials. We tested this by correlating the trial-wise decoding strengths of the cued and the rotated item (excluding no-rotation trials). A negative correlation would be evidence that only one of them was coded in any one trial, while the simultaneous maintenance of both items in each trial would result in a positive correlation due to variable noise levels of each trial. While it did not reach statistical significance, there was a trend of a positive correlation between the decoding strengths of the cued and the rotated item (*p* = 0.084, BF_10_ = 1.087; Fig. 6). Explicitly testing whether the correlation was negative provided strong evidence against it (BF_10_ = 0.074, one-sided). This provides evidence that there was no trade-off in the neural strengths of the items across trials and suggests that both items may have been present simultaneously in the neural data in at least some trials, which best fits with scenario 4Di above.

**Figure 6.**
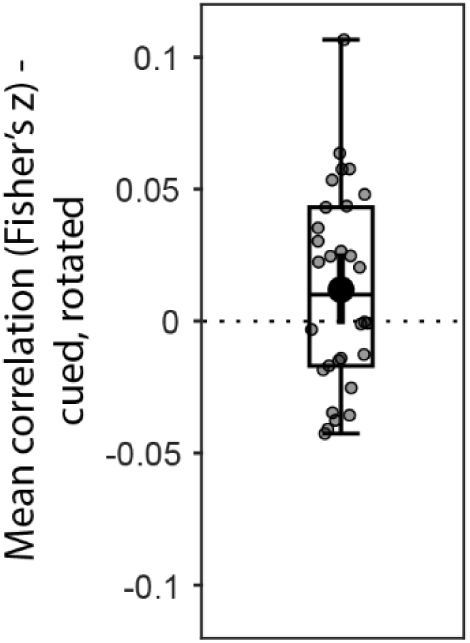
Mean correlation (Fisher’s z transformed) between the trial-wise decoding strengths of the cued/original item and the rotated item after transformation at Impulse 2. Error bars are the 95% C.I. of the mean.

#### 3.2.5 Generalization of coding schemes before and after transformation

We tested if the coding scheme used for the cued item before rotation (Impulse 1) generalized to the coding scheme after rotation (Impulse 2). Training the classifier on the cued item at Impulse 1 resulted in significant cross-generalization when tested on the cued item at Impulse 2 of rotation trials (*p* = 0.028, BF_10_ = 2.648; Fig. 7, left). This is evidence that even in the face of mental manipulation, the code of the original, non-manipulated item persists, and its coding scheme remains relatively stable over time. We also tested if the classifier trained on the cued item at Impulse 1 generalised to the mentally rotated item at Impulse 2. We found no evidence for this (*p* = 0.726, BF_10_ = 0.209; Fig. 7, right). The difference between the two was not significant (*p* = 0.204, BF_10_ = 0.318), however. This again suggests that the mental rotation of the cued item resulted in a new coding scheme for the rotated item, without removing the original item from the WM network, which remained relatively stable.

**Figure 7.**
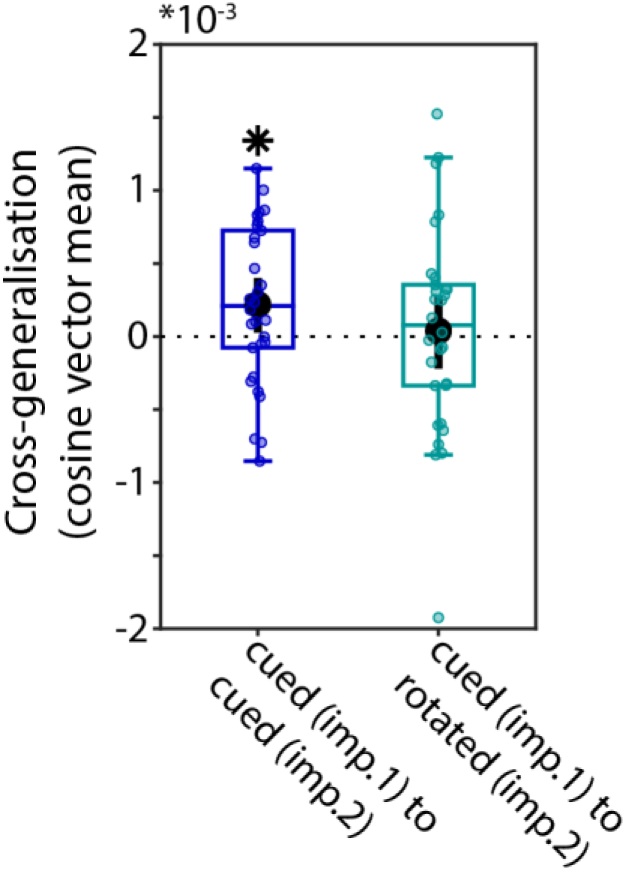
Cross-generalization of coding schemes between the original item before transformation at Impulse 1 and the original item and the rotated item after transformation at Impulse 2. The “0 rotation” condition is excluded. Error bars are 95% C.I. Statistically significant (*p* < 0.05) cross-generalization is marked with an asterisk (*).

#### 3.2.6 Relationship between trial-wise decoding strengths and WM performance

We tested if the trial-wise decoding strengths of the cued/original and the rotated item after transformation at Impulse 2 predicted trial-wise fluctuations in performance. Using logistic regression, we found no evidence that accuracy was predicted by either the cued/original item or the rotated item on a trial-by-trial basis (cued: *p* = 0.904, BF_10_ = 0.197; rotated: *p* = 0.197, BF_10_ = 0.555, Fig. 8A. Trial-wise variance in log-RT was predicted by both using linear regression, however (cued: *p* = 0.044, BF_10_ = 1.57; rotated: *p* = 0.021, BF_10_ = 4.809, Fig. 8B).

**Figure 8.**
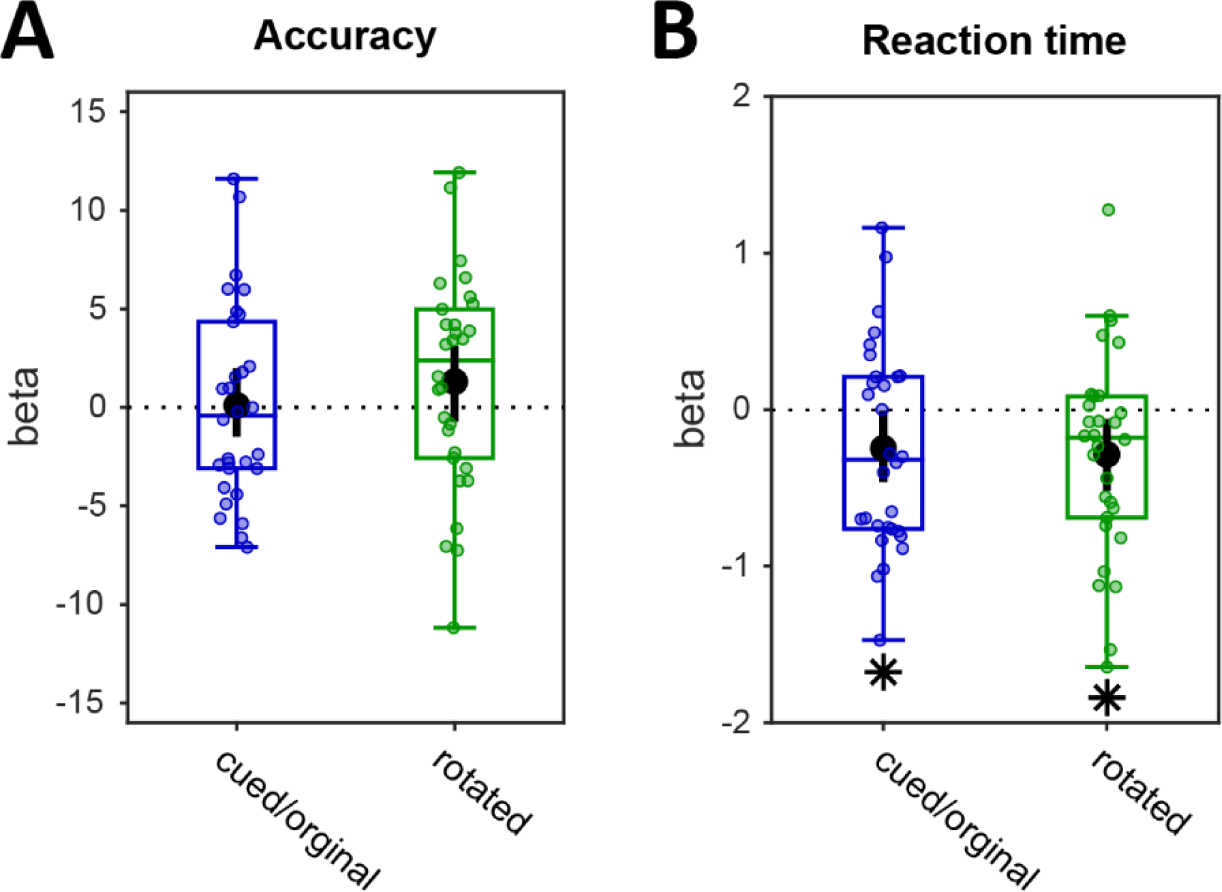
Relationship between trial-wise decoding at Impulse 2 and performance. **(A)** Regression weights (beta) of logistic regression between item decoding and accuracy. **(B)** Regression weights (beta) of linear regression between item decoding and log-transformed reaction times. The “0 rotation” condition is excluded. Error bars are 95% C.I.. **\*** *p* < 0.05.

## 4. Discussion

For working memory to guide behaviour in the flexible manner required by the dynamic environment we live in, it cannot rely on the storage and static maintenance of sensory input alone, but also needs to be able to manipulate and update information when necessary. Here, we assessed how working memory represents items that were not directly formed by sensory input, but were imagined by variable degrees of mental rotation. From the EEG response to an impulse stimulus presented during WM maintenance we were able to decode not only visually presented items, but also imaginary ones. We discovered that the originally presented memory items, although rendered task-irrelevant after mental rotation, continued to be represented concurrently with, and were coded independently from, the imagined rotation products. By contrast, uncued items seemed to be purged rapidly from WM, and were not decodable, as was previously observed (Wolff et al., 2017).

### 4.1 Representational coding of veridical and imagined items

As predicted by synaptic theories of WM (Mongillo et al., 2008; Zucker & Regehr, 2002), the visual impulse allowed an external readout of WM contents in the present study. Previous studies that have employed the impulse perturbation technique have shown its efficacy for stimuli that are presented in both the visual and auditory modality, and which are encoded in WM (Wolff et al., 2017; 2020a; 2020b). Here we found that the impulse is also effective for imagined items, supporting the idea that these items are similarly maintained in WM, and that they may equally utilise activity-quiescent brain states. The results furthermore suggest that the recovery of information in WM by means of a visual impulse is not dependent on the encoding of previous, direct sensory input, extending the scope of this approach. At the same time, the EEG patterns associated with the veridical (cued) item and the rotated item showed that their representations might differ in certain ways.

Although the cued item was clearly retained throughout the trial, change was observed in its neural representation across different time points. At the second impulse, the interval during which the cued item was significant was comparatively late, compared to the first impulse (236 to 428 ms post-impulse vs. 92 ms to 300 ms, respectively). Speculatively, this might be related to the change in task-relevance of the cued item at these timepoints. Furthermore, at Impulse 2, the representation of the cued orientation was no longer spatially specific. Earlier studies using multiple impulse signals reported that memory content remained spatially specific in time (Wolff et al., 2020b), which suggests that the loss of spatial specificity could be related to the present transformations. In a recent animal study, Panichello and Buschman (2021) observed that spatially specific memory content was transformed to another state with the involvement of the prefrontal cortex, once this memory representation became relevant for behaviour, and that this representation no longer carried spatial information (cf. Ester et al., 2009; Fukuda et al., 2016; Stokes et al., 2013). Likewise, in the current study, the cued orientation might have been transformed to a new state once it was relevant for mental rotation, and in which spatial information would not have a functional role. At the same time, we did observe cross-generalization of the cued item at Impulse 2 with that of Impulse 1, before rotation, suggesting that the coding scheme is otherwise still similar. Future research might clarify the role of transformations in the observed loss of spatial specificity by adding a systematic manipulation of the task-relevance of spatial information.

The results furthermore provided evidence that veridical and imagined items share the same representational substrate in visual processing areas of the brain, as both were sensitive to the visual impulse signal, and decodable from posterior electrodes alone. However, we also observed that there was no representational similarity between the original, cued orientation and the rotation product at the second impulse. That is, a given orientation angle (e.g., 30°) was represented differently, depending on whether it was previously presented on the screen, or the end product of mental rotation (e.g., 60° rotated 30° counter-clockwise). This finding seems to contradict those of the fMRI study by Stokes and colleagues (2009) on mental imagery, in which mental imagery and visual perception activated representations in the same areas of the visual cortex. Similar correspondence was later reported for items held in WM (Albers et al., 2013), and Christophel and colleagues (2015) also observed that original and rotated colour patches seemed to share a similar coding scheme in visual and parietal cortices. It is possible that the discrepancy between the currently observed lack of representational similarity between rotated and original items, and the similarity observed by the aforementioned authors is due to the inherent temporal blurring of fMRI as well as their analysis approach, in which multiple time points were combined. It is also possible that subtle differences in the level of abstraction may have led to differences in coding scheme, as has been reported for auditory stimuli before (Linke & Cusack, 2015). Nevertheless, our results also show important commonalities between rotated and original items in that both were revealed in the impulse response, which is in line with these studies.

The representational difference between the original and rotated item that we observed contradicts a strict interpretation of memory models that define working memory as an activated and attended portion of long-term memory (e.g., Cowan, 1999; 2005; Oberauer, 2002; 2009). In such models, a single memory system houses all representations. The representations only differ with regard to their activity level; high levels correspond to items held in WM and in the focus of attention, while low levels correspond to items held in long-term memory. Accordingly, since in our design the visually presented items as well as the rotation products consisted of the same 6 orientation angles, these representations should have cross-generalized: Mental rotation should have reactivated the same, shared orientation representations from the low activity state as visually presented items would have. As indicated, our results showed instead that the veridical and imagined representations did not cross-generalize. Since veridical items did generalize over time, from the first to the second impulse, despite a loss of spatial specificity, the lack of cross-generalization between veridical and transformed representations cannot be ascribed to a lack of power. More generally, this lack of cross-generalization also supports the idea that our participants performed mental rotation by actually generating new images, rather than recalling discrete items from long-term memory.

Diverging from our results, a recent study showed cross-generalization of maintained and transformed WM representations of spatial locations (Günseli et al., 2024). In this study, participants were asked to maintain a location, followed by a transformation of spatial locations. Maintained and transformed spatial WM representations shared a common representational format, as measured by cross-generalization of alpha activity. The relationship that exists between alpha activity and covert attention is one possible explanation for these diverging results, as alpha-band activity tracks the locus of covert attention (Kelly et al., 2006; Thut et al., 2006; Worden et al., 2000). Therefore, the cross-generalization between maintained and transformed spatial WM representations in Günseli et al. (2024) could be explained by covert attention moving from one location (the encoded location) to another (the transformed location).

### 4.2 Capacity limits

In view of the scarcity of WM space, possibly the most striking outcome of the present study was the novel observation that obsolete original items were retained in WM concurrently with the task-relevant rotation products that were derived from them. It is conceivable that this liberal use of WM space is also one of the factors that make mental rotation a relatively difficult task. The rotation process itself is certainly not trivial, as was evident from the progressive reduction in performance for increasing angles of rotation that we presently observed, and which is commonly found in mental rotation paradigms (e.g., Wexler et al., 1998; Searle & Hamm, 2017). After the rotation itself, the way in which both the now-irrelevant starting point and the outcome of this transformation process are maintained in WM may further compound that difficulty.

By contrast, previous work has shown that items that are no longer task-relevant, such as those that are uncued, are quickly purged, and can no longer be decoded even after impulse perturbation (Wolff et al., 2017). This has supported the view that WM may employ an active purging mechanism to get rid of information that is no longer needed. Alternatively, though, such information may also simply fade from WM, due to its inherently volatile nature, in the absence of active reinforcement (e.g., periodic refreshing). The latter account is in line with predictions from a recent computational model of WM, based on calcium-mediated short-term synaptic plasticity (Pals et al., 2020). The current results nevertheless provide further evidence that a purging mechanism may indeed exist, since the fate of irrelevant items, that is, uncued items and original items after rotation, was not the same. This may indicate that the brain treated them differently, such that the former, but not the latter items were actively removed.

There may be a good reason why the brain seems to maintain the obsolete original items. In the current task, holding onto the original item as well as the rotation instructions allows the participants to recreate or double-check the rotation product. There was some evidence that this helped to improve task performance; lower probe response times were associated with better decoding of both original and rotated items. While a memory-related account of this effect is most parsimonious, it should be noted that the impulse response might also be mediated by attentional processes. Nevertheless, in everyday scenarios it may also often make sense to remember more than the end-product of a mental transformation. For instance, if we predict the future location of a temporarily occluded vehicle in the environment, it would be useful to do so flexibly, to be able to make use of different estimates of its speed. Such flexibility requires the retention of the original input (the location of the vehicle) and the transformation (estimated distance covered based on speed), so that they can be used again and adjusted as needed. Thus, we propose that the maintenance behaviour observed in our experiment might reflect the prioritization of adaptive flexibility over WM storage capacity, despite the scarcity of the latter. Whether the retention of the original input is an active process, or the result of more passive mechanisms that leave the original item in place, remains to be determined.

In this context it may be worth noting that in the study of WM, a lot of research has been devoted to charting WM capacity limits: The number of items (e.g., Miller, 1956; Cowan, 2001), the organization of information in individual properties and compound objects (e.g., Luck & Vogel, 1997; Xu, 2002), and the nature of WM capacity itself in terms of continuous resources or discrete slots (e.g., Zhang & Luck, 2008; Bays et al., 2009). The insights gained from this long-standing and important line of research remain highly relevant to this date. However, the present work suggests that a full understanding of WM cannot reflect on storage capacity alone. It also needs to develop a perspective on how the available storage capacity in WM may be utilized to support adaptive behaviour. The current results suggest that at times, the already strongly limited capacity of WM is filled very rapidly. From a strict capacity perspective, there would be little reason to assume that WM load at any one time after the initial retro-cue in our experiment would be more than a single item, yet the data showed differently. It is crucial to further our understanding of WM by examining the conditions that may foster such capacity-costly behaviour.

### 4.3 Conclusion

In line with synaptic theories of WM, we found that the representations of mentally rotated items and of visually presented items rely on the same neural substrate, as both could be decoded during maintenance from the EEG impulse response, even though their coding schemes appeared to be different. Importantly, in doing so the brain seems willing to sacrifice already-scarce WM capacity to support flexible behaviour in dynamic environments, as we observed that the original, no-longer task-relevant items continued to be maintained concurrently with transformed ones. This finding prompts the question of how much WM capacity is commonly ‘lost’ by this striking tendency to hold on to obsoleted information.

## Author Contributions

Güven Kandemir: Conceptualization, Methodology, Formal analysis, Investigation, Data curation, Writing – original draft, Visualization.

Michael J. Wolff: Methodology, Software, Validation, Formal analysis, Data curation, Writing –review & editing, Visualization.

Aytaç Karabay: Conceptualization, Methodology, Validation, Investigation, Writing –review & editing Mark G. Stokes: Conceptualization, Methodology, Supervision, Project administration, Funding acquisition

Nikolai Axmacher: Conceptualization, Writing –review & editing, Supervision, Project administration, Funding acquisition

Elkan G. Akyürek: Conceptualization, Methodology, Writing –original draft, Writing –review & editing, Supervision, Project administration, Resources, Funding acquisition.

## Supplementary Materials

**Supplemental Figure 1.**
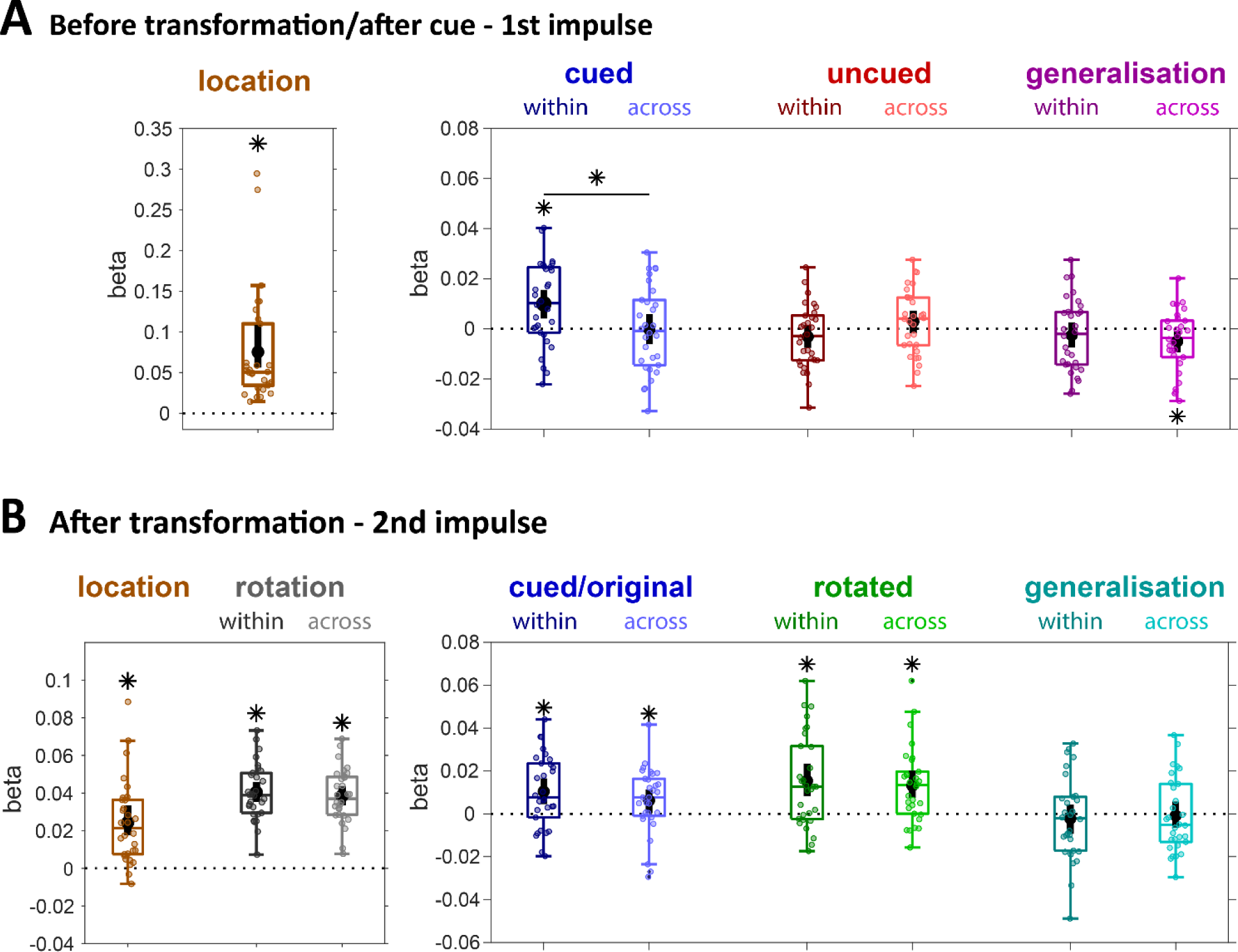
LDC results of the spatiotemporal signal using a pre-trial baseline (-200 to 0 ms, relative to memory onset). **A.** Beta values of model fits for Impulse 1. Cued location model: *p* < 0.01, BF_10_ > 1000. Cued items models: within location, *p* < 0.01, BF_10_ = 39.958; across location, *p* = 0.974, BF_10_ = 0.196, difference; *p* = 0.018, BF_10_ = 4.395. Uncued item models: within, *p* = 0.172, BF_10_ = 0.453; across, *p* = 0.246, BF_10_ = 0.423; difference, *p* = 0.084, BF_10_ = 1.365. Generalisation models (cued/uncued): within, *p* = 0.338, BF_10_ = 0.304; across, *p* = 0.032, BF_10_ = 1.946; difference, *p* = 0.406, BF_10_ = 0.259. **B.** Beta values of model fits for Impulse 2. Cued location model: *p* < 0.01, BF_10_ > 1000. Rotation condition models: within location, *p* < 0.01, BF_10_ > 1000; across location, *p* < 0.01, BF_10_ > 1000, difference; *p* = 0.826, BF_10_ = 0.354. Cued items models: within location, *p* < 0.01, BF_10_ = 37.709; across location, *p* = 0.042, BF_10_ = 1.655, difference; *p* = 0.456, BF_10_ = 0.258. Rotated item models: within, *p* < 0.01, BF_10_ = 85.023; across, *p* < 0.01, BF_10_ = 201.893; difference, *p* = 0.878, BF_10_ = 0.213. Generalisation models (cued/rotated): within, *p* = 0.548, BF_10_ = 0.255; across, *p* = 0.824, BF_10_ = 0.2; difference, *p* = 0.744, BF_10_ = 0.199.

**Supplemental Figure 2.**
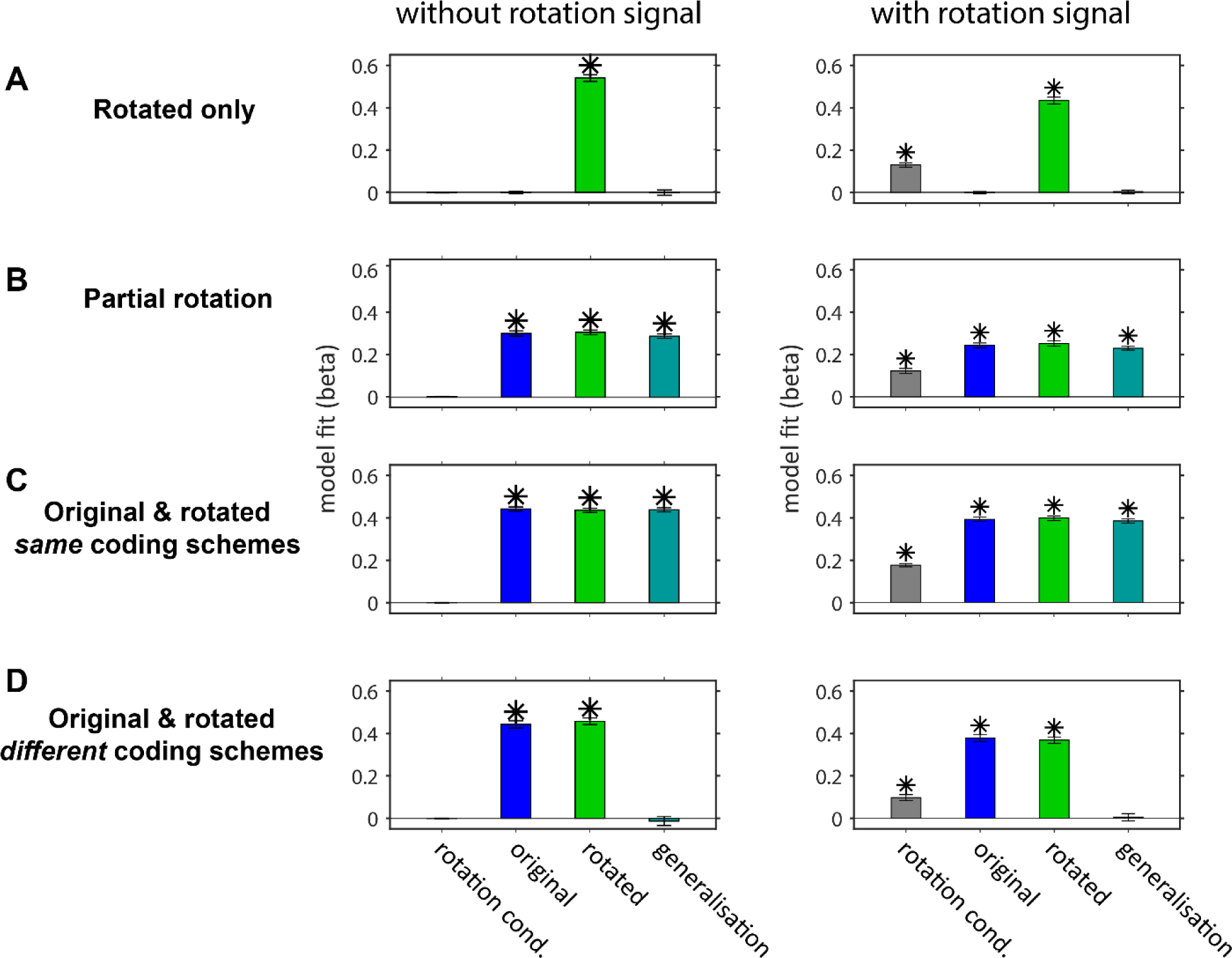
Simulation results when an explicit rotation signal was not added (left column, as in Fig. 4), and when it was added (right column).

**Supplemental Figure 3.**
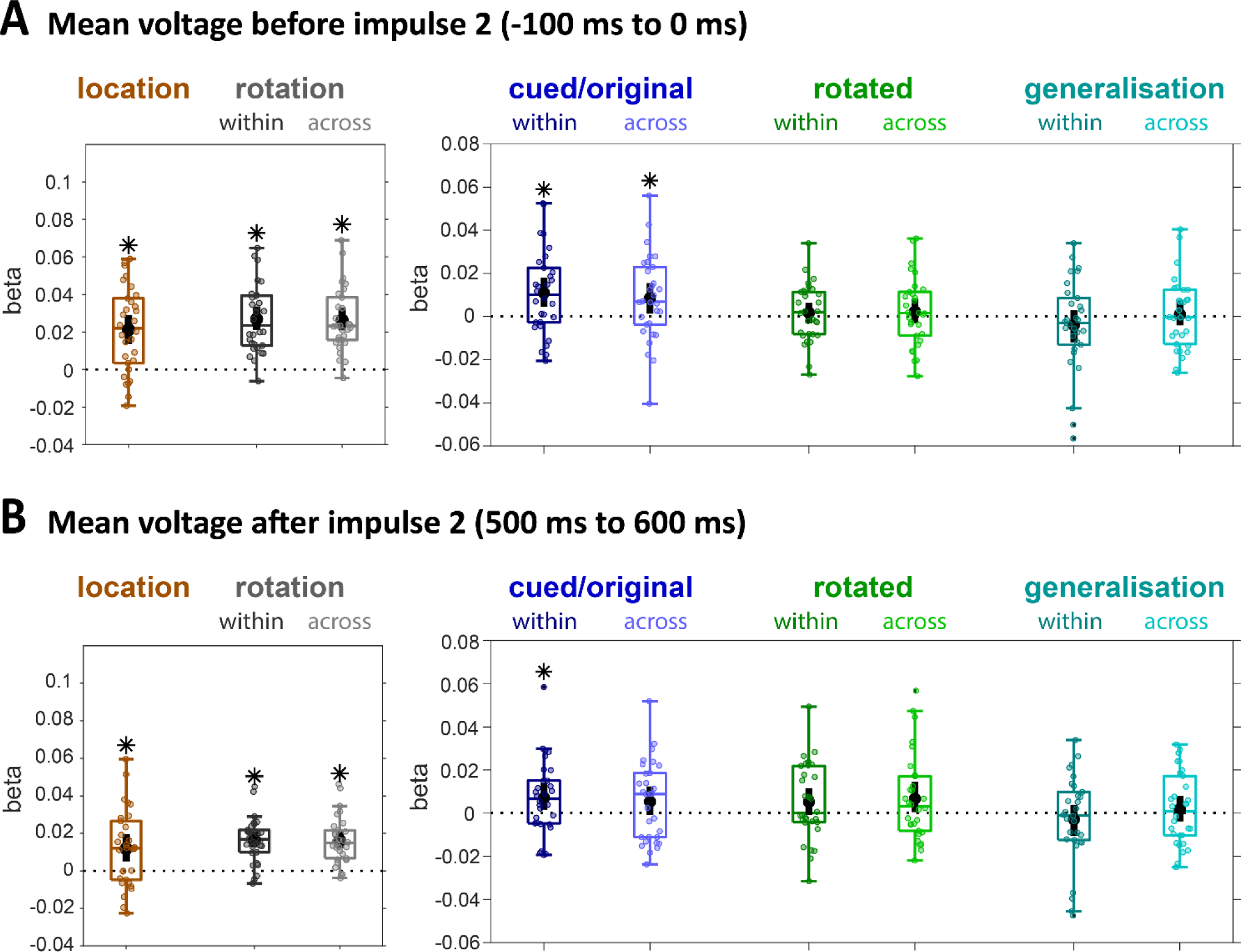
LDC results at Impulse 2 of the mean voltage immediately before (-100 ms to 0 ms relative to Impulse 2), and after Impulse 2 presentation (500 ms to 600 ms relative to Impulse 2 and -100 ms to 0 relative to Probe onset), using a pre-trial baseline (-200ms to 0, relative to memory array onset). **A.** Beta values of model fits before Impulse 2. Cued location model: *p* < 0.01, BF_10_ > 1000. Rotation condition models: within location, *p* < 0.01, BF_10_ > 1000; across location, *p* < 0.01, BF_10_ > 1000, difference; *p* = 0.876, BF_10_ = 0.204. Cued items models: within location, *p* < 0.01, BF_10_ = 17.248; across location, *p* = 0.03, BF_10_ = 4.179, difference; *p* = 0.636, BF_10_ = 0.201. Rotated item models: within, *p* = 0.448, BF_10_ = 0.272; across, *p* = 0.448, BF_10_ = 0.270; difference, *p* = 0.972, BF_10_ = 0.199. Generalisation models (cued/rotated): within, *p* = 0.312, BF_10_ = 0.314; across, *p* = 0.774, BF_10_ = 0.196; difference, *p* = 0.346, BF_10_ = 0.598. **B.** Beta values of model fits after Impulse 2. Cued location model: *p* < 0.01, BF_10_ = 24.482. Rotation condition models: within location, *p* < 0.01, BF_10_ > 1000; across location, *p* < 0.01, BF_10_ > 1000, difference; *p* = 0.692, BF_10_ = 0.204. Cued items models: within location, *p* = 0.044, BF_10_ = 3.259; across location, *p* = 0.126, BF_10_ = 0.763, difference; *p* = 0.616, BF_10_ = 0.208. Rotated item models: within, *p* = 0.098, BF_10_ = 0.565; across, *p* = 0.060, BF_10_ = 0.583; difference, *p* = 0.884, BF_10_ = 0.214. Generalisation models (cued/rotated): within, *p* = 0.448, BF_10_ = 0.248; across, *p* = 0.582, BF_10_ = 0.583; difference, *p* = 0.350, BF_10_ = 0.459.

**Supplemental Figure 4.**
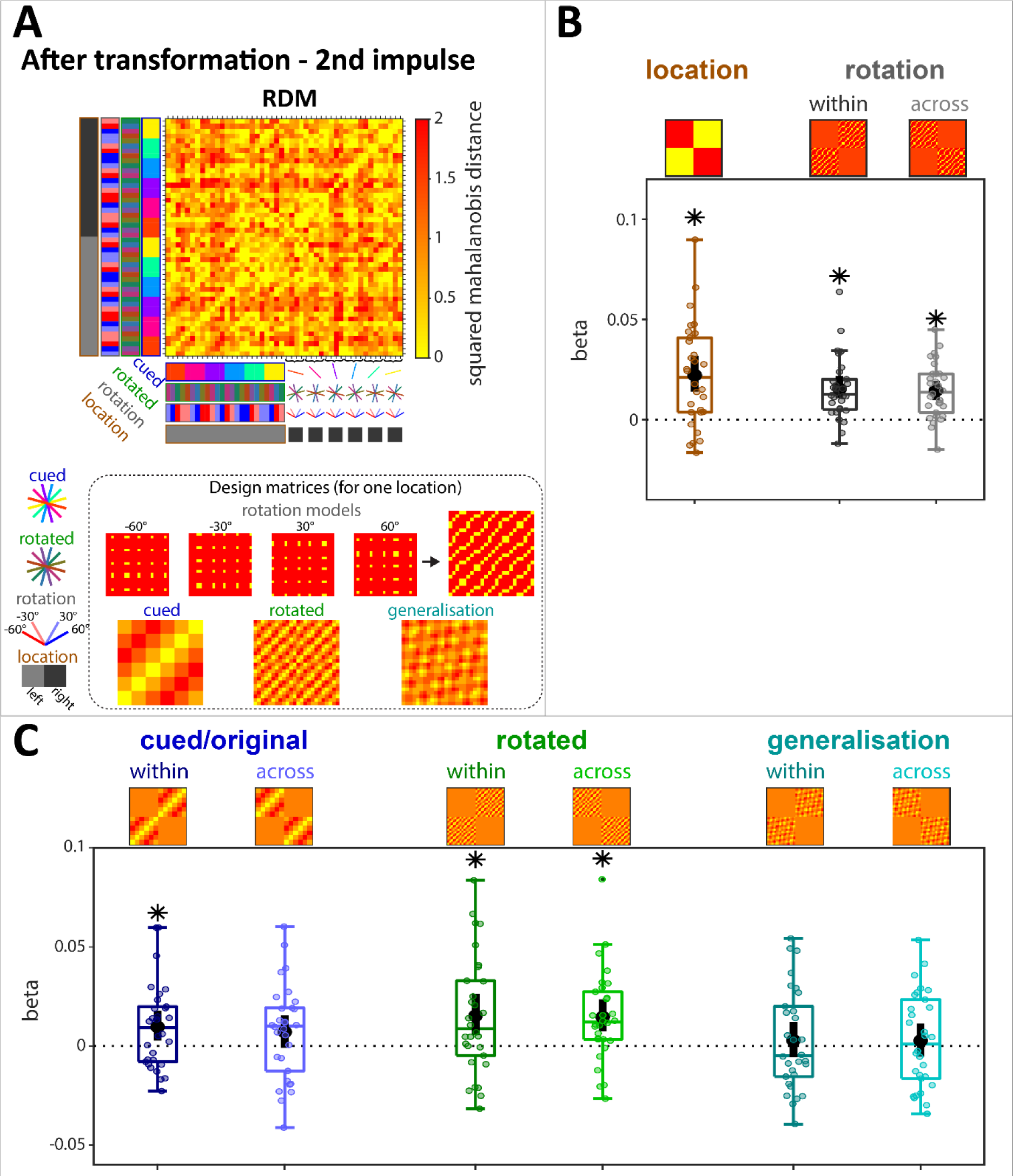
LDC of the spatiotemporal signal at Impulse 2 when excluding all non-rotation trials. **A.** Average RDM and design matrices of models. **B.** Beta values of the cued location model (*p* < 0.01, BF_10_ = 582.24) and the rotation condition models (within and across location: *p* < 0.01, BF_10_ > 1000; difference: *p* = 0.784, BF_10_ = 0.218). **C.** Beta values of cued item models (within: *p* = 0.026, BF_10_ = 4.502; across: *p* = 0.14, BF_10_ = 0.571; difference: *p* = 0.51, BF_10_ = 0.208), rotated item models (within: *p* = 0.014, BF_10_ = 7.339; across: *p* < 0.01, BF_10_ = 132.813; difference: *p* = 0.652, BF_10_ = 0.196), and generalization models (within: *p* = 0.538, BF_10_ = 0.239; across: *p* = 0.538, BF_10_ = 0.265; difference: *p* = 0.986, BF_10_ = 0.196).

**Supplemental Figure 5.**
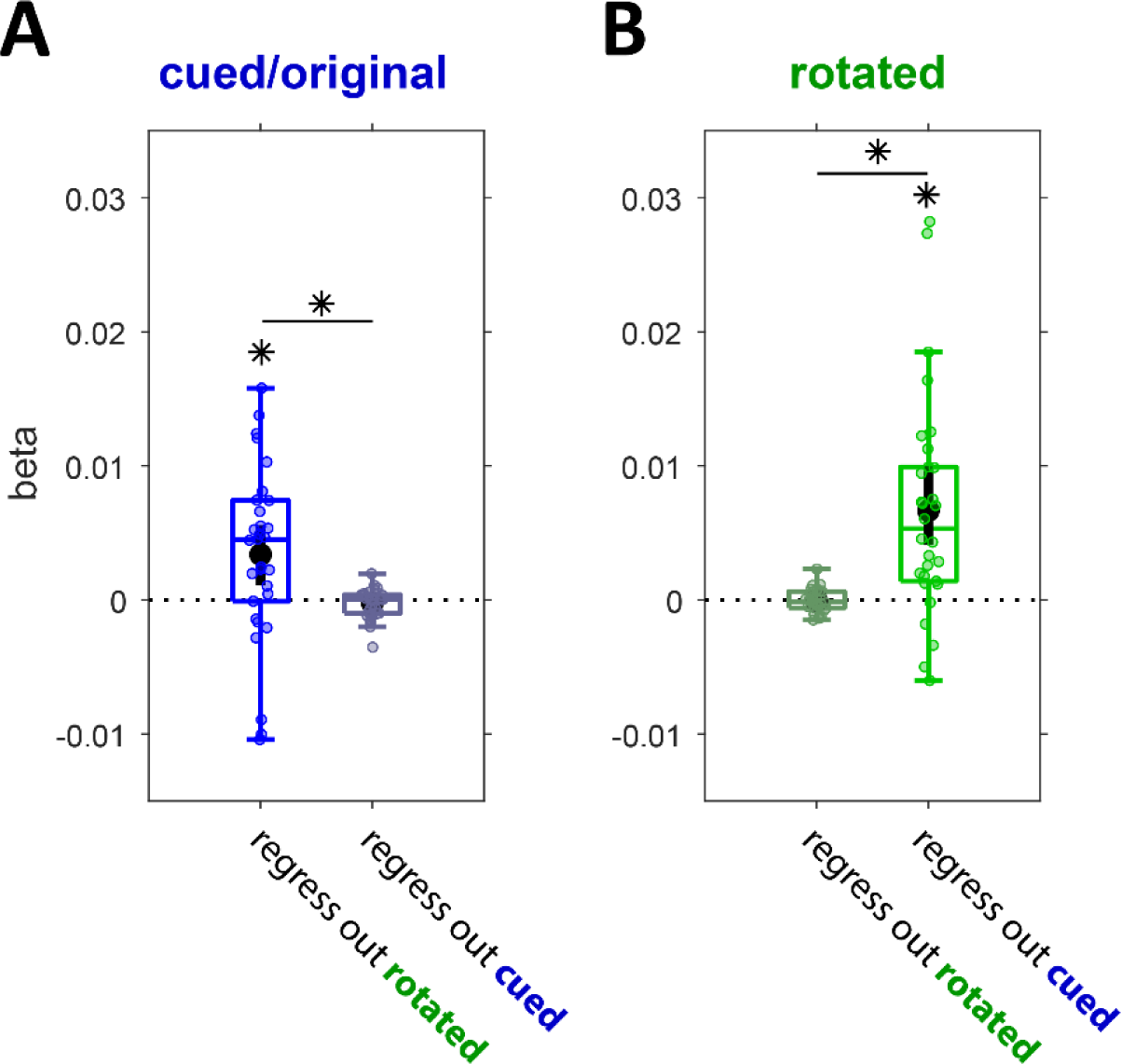
Model fits of cued and rotated item models at Impulse 2 when the effect of one a specific model (cued or rotated item) is estimated from an independent half of the data, and then regressed out from the other half. **A.** Cued item model fit when independently regressing out the rotated item effect (*p* < 0.01, BF_10_ = 12.572), and the cued item effect (*p* = 0.21, BF_10_ = 0.312). Difference: *p* < 0.01, BF_10_ = 17.094). **B.** Rotated item model fit when independently regressing out the rotated item effect (*p* = 0.7686, BF_10_ = 213), and the cued item effect (*p* < 0.01, BF_10_ = 856.518). Difference: *p* < 0.01, BF_10_ > 1000).

